# A Vision-Language Model as a Teacher for Bird Vocalization Detection

**DOI:** 10.64898/2026.09.23.753955

**Authors:** George Vengrovski, Timothy J. Gardner

## Abstract

Time-frequency detection of bird vocalizations is an important step toward turning weakly annotated field recordings into usable data for studying avian communication. Supervised detectors trained on human expert annotations scale poorly across species and recording conditions, so we introduce a teacher-student setup in which the teacher, a vision-language model, labels bounding boxes on spectrograms of citizen-scientist recordings, and those labels train a student, a self-supervised bioacoustic encoder. We find that the student generally exceeds both the teacher and supervised models trained on human annotations and performs strongly on held-out datasets for both time-frequency and onset-offset localization. YOLO detectors trained on our teacher labels match those trained on human annotations, suggesting that VLM labels can substitute for costly expert labeling.

## 1. Introduction

Deep learning has turbocharged the study of avian communication, allowing researchers to automate previously labor-intensive tasks. In particular, acoustic encoders trained on large amounts of recordings have led to success in classifying species in recordings [1], detecting the presence of bird vocalizations in recordings [2], and even the autonomous discovery of units of song [3]. This has enabled the monitoring of bird populations [4], migratory patterns [5], and even song structure analysis in neuroscience experiments [6].

Much of this progress has relied on large datasets from citizen scientists [7], who, using mobile phones or low-cost autonomous recording devices, have collected diverse data world-wide. However, these recordings are usually only weakly annotated, often with a single species tag for the entire file, even though they typically contain long stretches of background noise, vocalizations from multiple individuals and species, and anthropogenic sounds (such as airplanes).

A method that effectively localizes bird vocalizations in time and frequency would help turn these weakly annotated recordings into usable data for studying avian communication [8]. Because audio is typically represented as a spectrogram, an image with time and frequency axes, this localization can be framed as bounding-box detection in images. Temporal boundaries enable measurements of vocal duration and timing, while frequency boundaries allow analyses to exclude sounds outside the vocalization’s frequency range, including some overlapping calls and background noise. These localizations could also support self-supervised learning by directing training toward vocal regions rather than unrelated background sounds, while reducing the computational cost of training and inference for bioacoustic models by excluding background noise and non-avian vocalizations.

Despite the importance of isolating these bird vocalizations, detectors that work across species and recording conditions remain in their infancy, with substantial progress only recently made [9, 10]. Traditional supervised learning methods are expensive and labor-intensive because they require many annotators familiar with birdsong to annotate across different locations and species. Producing thousands of such annotations is therefore expensive and difficult to scale. Moreover, as new recording environments and devices appear, additional human annotation may be required to maintain performance under the resulting distribution shifts.

Modern vision-language models (VLMs) provide a possible alternative to large-scale human annotation. We use a VLM to produce a substantial amount of bounding-box annotations across many bird species, locations, and conditions collected from citizen scientists (Fig. 1). Powerful VLM models suitable for such an annotation task are large and slow, so we distill the produced time-frequency annotations as supervised teacher labels for an existing self-supervised bioacoustic encoder, “Song-MAE”, which results in an improvement in annotation over the VLM teacher and achieves a high level of performance in bird vocalization detection.

**Fig. 1.**
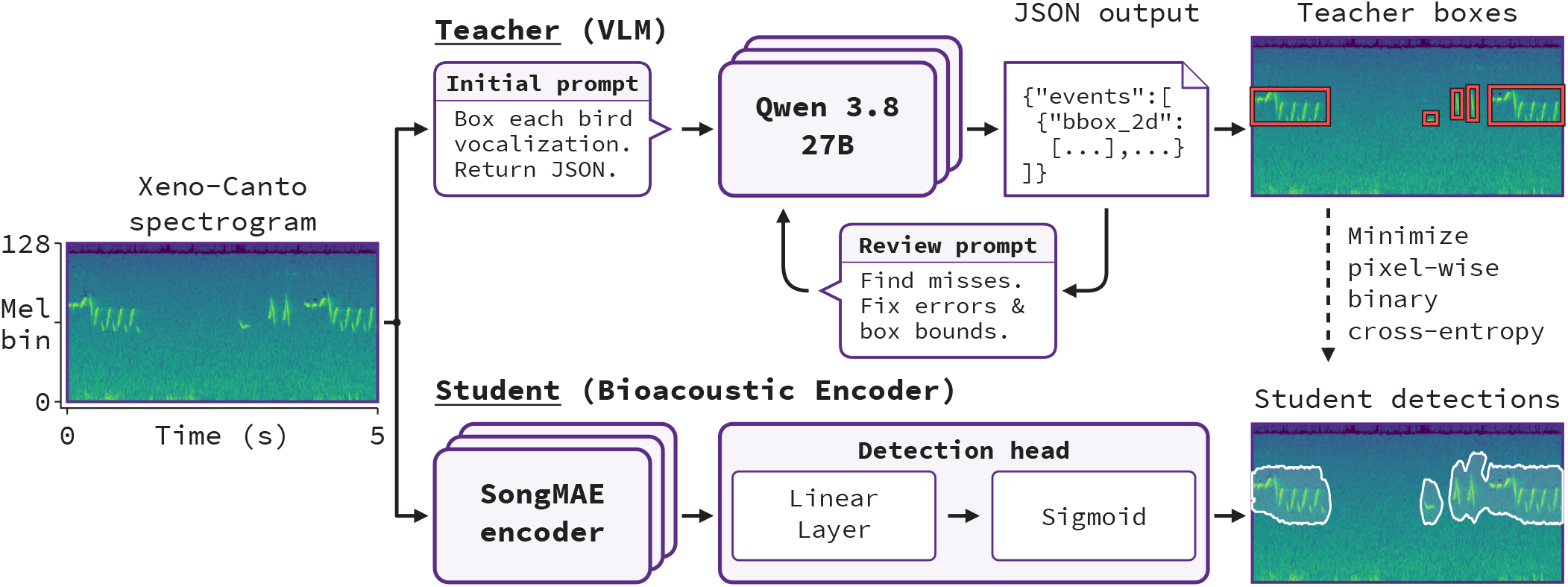
Overview of the pipeline. A VLM teacher (Qwen3.8-27B) annotates Xeno-Canto spectrograms with bounding boxes through an initial prompt and a self-review pass. These boxes supervise a student bioacoustic encoder (SongMAE) with an additional detection head.

## 2. Related Works

Large-scale bioacoustic modeling has primarily focused on clip-level audio tagging and species classification. Models such as Bird-MAE and BirdAVES (a bird-specific release of AVES) are pretrained on large collections of bird recordings and produce general-purpose representations that transfer to both tasks [11, 12].

Several methods have focused on directly extracting fore-ground bird vocalizations. Bambird combines time-frequency segmentation with unsupervised clustering to produce spatiotemporal localizations of birdsong [8]. Another approach detects song by denoising birdsong spectrograms [13]. More recently, YOLO (You Only Look Once) [14, 15], an object-detection architecture that predicts bounding boxes around objects in images, has been utilized to place bounding boxes around bird vocalizations [10] in tropical soundscapes. Although effective, these approaches rely on human-produced labels and are trained and evaluated in narrow contexts. BirdCODE, however, is trained using only weak human labels, via synthetic mixtures and pseudolabels, and produces temporal onset and offset annotations without frequency localization [9].

Separately, VLM models have been shown to produce few-shot classification of audio spectrograms [16]. We have not found any studies that show that VLM models can localize bird vocalizations.

## 3. Methods

### 3.1 VLM Teacher

For the VLM teacher, we chose Qwen3.8-27B, a dense 27-billion-parameter vision-language model, for its strong visual reasoning and agentic performance, while remaining small enough to run locally on consumer-grade hardware [17]. We served Q8 0-quantized weights locally through llama.cpp.^1^

For generating annotations, we first computed a mel spectrogram for each source recording (32 kHz mono, 1024-sample Hann window, 5 ms hop, 128 mel bins, power scaled to decibels relative to the recording maximum) and then split it into non-overlapping 5-second windows, zero-padding any window shorter than 5 seconds. We then rendered each window as a 2048 × 512 viridis PNG (2240 × 704 with coordinate axes), choosing this high resolution so that the VLM would allocate more vision tokens to each spectrogram. We passed each image to the VLM with a system prompt instructing it to tightly localize avian vocalizations and return bounding box coordinates and confidence scores in JSON format. We normalized the coordinates to the range 0–1000 relative to the spectrogram rectangle.

We built the VLM teacher annotation pipeline as a series of stages (Fig. 1), which we later ablate (Sec. 5.1) to determine which stages improve annotation quality. The stages are as follows:

#### 1. Initial Prediction

We pass the prompt and spectrogram image to the VLM, and the VLM teacher outputs annotations. As ablations, we also run this stage with chain-of-thought (CoT) reasoning disabled and test whether providing explicit frequency and time coordinate axes benefits annotation.

#### 2. Self-review

The VLM re-inspects the clean spectrogram alongside a second image with the initial boxes rendered in red and numbered, along with the previous detections as JSON, and returns a complete revised detection list that corrects missed events, false positives, and inaccurate boundaries.

#### 3. Second self-review

We apply the same review instruction again to the output of the first self-review.

### 3.2 SongMAE Student

For the student model, we use SongMAE, a self-supervised vision transformer pretrained on a diverse set of recordings containing bird vocalizations [18]. SongMAE is well suited to this task because it encodes a spectrogram as a fine grid of latents spanning both time and frequency, with each latent covering 32 mel bins and 5 ms. We fine-tune the SongMAE encoder and train a linear detection head on top of it. For each latent, the head outputs 32 logits, one per mel bin, which we pass through a sigmoid to obtain the probability that each time-frequency bin belongs to a bird vocalization.

We train the backbone and detection head jointly with binary cross-entropy against masks derived from the VLM teacher boxes, where every time-frequency bin inside a box is labeled as bird vocalization and all others as background. We optimize for 5 epochs with AdamW [19], using a learning rate of 10*−*5 for the backbone and 10*−*3 for the head, a weight decay of 10*−*4, and a batch size of 16, and keep the epoch with the lowest validation loss for evaluation. At inference, we Gaussian-smooth the fore-ground probability map with standard deviations of 2 mel bins in the frequency domain and 3 frames (15 ms) in the time domain, then binarize it using a single threshold.

### 3.3 Evaluation Metrics

Although our primary goal is time-frequency localization of bird vocalizations, we also evaluate onset-offset localization as a secondary task, since it enables comparison with the BirdCODE model and is sufficient for some downstream analyses, such as measuring vocalization durations. For both tasks, we report average precision (AP) as our primary metric and mean intersection over union (IoU) as a secondary metric. AP is threshold-free and measures ranking quality, whereas IoU scores a single thresholded output, matching how detectors like YOLO are deployed. We select this threshold on the Powdermill development set and hold it fixed for all evaluation. For onset-offset evaluation, we collapse the 2D outputs of YOLO11 and our model along the frequency dimension. Since the temporal resolution of the models we compare differs, we rasterize all predictions and annotations onto a common 5 ms grid.

## 4. EXPERIMENTS

### 4.1 Datasets and Splits

For the teacher-labeled training set, we use recordings from Xeno-Canto, organized in the BirdSet dataset [20], and annotate 5,500 unique recordings with the VLM teacher model. This yields 25,000 seconds of training data and 2,500 seconds of validation data, containing 31,422 unique labeled events from 5,500 species and 4,969 recording sites.

For the development set, we use the Powdermill dataset [21] (Fig. 2), which contains strong human annotations of bird vocalizations in both temporal and frequency extent. We use this set to select segmentation thresholds, ablate the VLM teacher pipeline, and examine the size of the SongMAE audio encoder backbone and the number of teacher labels. We conduct the final evaluation on the held-out XC-AJ [22], WABAD [23], NIPS4Bplus [24], and Hawaii [25] test sets. We use the WABAD and Hawaii datasets for two-dimensional localization, and XC-AJ and NIPS4Bplus for onset-offset localization.

**Fig. 2.**
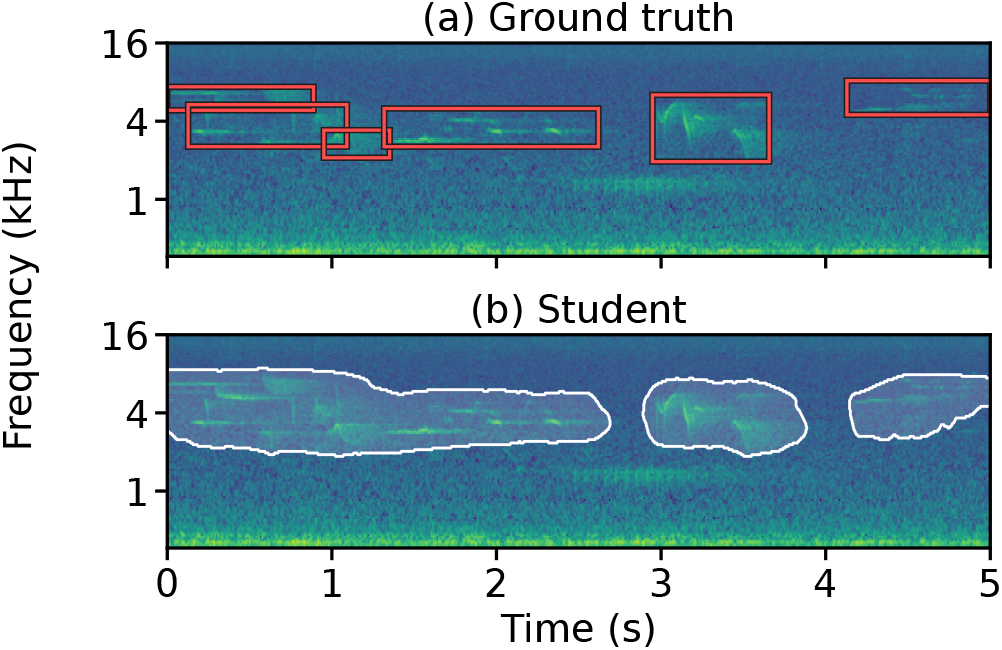
Powdermill development set human bounding-box annotations (a) and SongMAE-Large student prediction (b) for the same recording excerpt.

### 4.2 Model Comparisons

For time-frequency localization, we compare our SongMAE student against the released BirdBox YOLO11 nano and YOLO11 large detectors [10] (hereafter YOLO11n and YOLO11l), as well as YOLO11n and YOLO11l models that we train from COCO-pretrained weights on the same VLM teacher labels as Song-MAE, following the training procedure of [10]. This allows us to separate the effect of the teacher annotations from the effect of the student architecture. We also compare three SongMAE encoder sizes—Micro (1.75M parameters), Base (14.9M parameters), and Large (98.7M parameters)—using the same training and post-processing procedures for each. For every model we train using the teacher labels, we train it with three random seeds and report the mean. For one-dimensional temporal localization, we compare against BirdCODE and the temporally collapsed outputs of SongMAE, YOLO11n, and YOLO11l.

## 5. RESULTS AND DISCUSSION

### 5.1 VLM Teacher and Student Ablation

We ablate each stage of the teacher pipeline on the Powdermill development set, adding stages cumulatively (Table 1). Providing coordinate axes on the rendered spectrogram and enabling CoT reasoning give the largest gains, raising pixel AP from 0.242 to 0.452. Self-review adds a smaller improvement. Qualitatively, predictions without self-review contain errors such as shifted boxes or boxes placed on sounds that are not bird vocalizations, which review typically corrects; however, these affect only a small fraction of events and move the aggregate metrics only slightly. A second self-review pass gives only a negligible further gain, so we annotate the training set with a single initial prediction followed by one self-review.

**Table 1.** Effect of teacher annotation stages, added cumulatively, on localization performance on the Powdermill development set.

| VLM annotation pipeline | pixel AP |
| --- | --- |
| Initial prediction | 0.242 |
| Coordinate axes | 0.308 |
| CoT reasoning | 0.452 |
| Self-review | 0.457 |
| Additional self-review | **0.458** |

On the Powdermill development set, student performance improves with the amount of teacher-labeled training data, but the amount of data needed to reach high performance depends on backbone size (Fig. 3). SongMAE-Large reaches a pixel AP of 0.760 with only 1,000 seconds of training data, SongMAE-Base at 2,500 seconds, and SongMAE-Micro only at 10,000 seconds. Beyond 10,000 seconds, all three backbones converge to a pixel AP of about 0.77. We therefore use SongMAE-Large trained on the full 25,000 seconds for all downstream evaluations. The convergence may reflect a performance ceiling on Powdermill, or it may indicate that the student is limited by the teacher labels rather than by its own capacity, since additional teacher-labeled recordings may contribute few new types of vocalization and repeat the same systematic teacher errors.

**Fig. 3.**
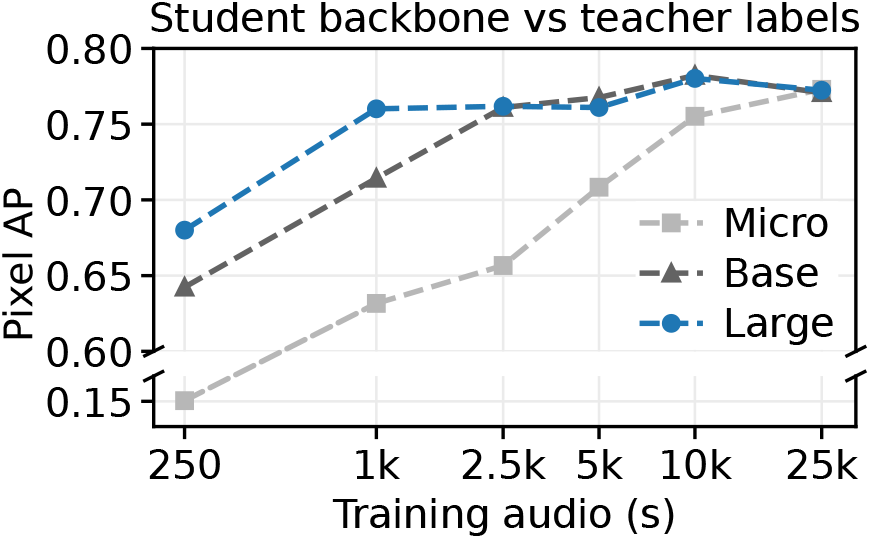
Student pixel AP on the Powdermill development set as a function of backbone size and amount of teacher-labeled training audio. Values are means across three seeds.

### 5.2 Time-frequency localization

On both held-out datasets, SongMAE-Large outperforms all YOLO11 detectors on pixel AP and 2D IoU, including those trained on our teacher labels (Table 2). The largest margin is on the Hawaii dataset, where pixel AP rises from 0.619 for the strongest YOLO11 model (YOLO11n trained on our teacher labels) to 0.716 for SongMAE. YOLO11 models trained on our teacher labels match or exceed the released models trained on human annotations (pixel AP 0.568 vs. 0.549 on WABAD and 0.604 vs. 0.546 on Hawaii for YOLO11l). This suggests that VLM teacher labels can substitute for human annotations when training a detector, and that SongMAE’s gains over YOLO11 stem from the student model rather than the labels. IoU gains are smaller than AP gains, suggesting that the binarization threshold calibrated on Powdermill does not fully transfer to the held-out datasets.

**Table 2.** Localization performance on the held-out test sets. WABAD and Hawaii (pixel-level AP and IoU) are evaluated for time-frequency localization, and XC-AJ and NIPS4Bplus (frame-level AP and IoU) for onset-offset localization. For models trained on our teacher labels, values are means across three seeds. Bold marks the best result in each column (ties share it), underline marks the second best.

| Model | WABAD |  | Hawaii |  | XC-AJ |  | NIPS4Bplus |  |
| --- | --- | --- | --- | --- | --- | --- | --- | --- |
|  | AP | IoU | AP | IoU | AP | IoU | AP | IoU |
| BirdCODE | — | — | — | — | **0.836** | 0.526 | 0.657 | **0.545** |
| YOLO11n (human labels) | 0.552 | 0.347 | 0.541 | 0.369 | 0.700 | <u>0.546</u> | 0.630 | 0.455 |
| YOLO11l (human labels) | 0.549 | 0.354 | 0.546 | 0.381 | 0.676 | **0.556** | 0.590 | 0.464 |
| YOLO11n (teacher labels) | 0.566 | 0.348 | <u>0.619</u> | <u>0.412</u> | 0.769 | 0.512 | <u>0.749</u> | 0.468 |
| YOLO11l (teacher labels) | <u>0.568</u> | <u>0.359</u> | 0.604 | 0.400 | <u>0.774</u> | 0.518 | 0.748 | <u>0.532</u> |
| SongMAE-Large (teacher labels) | **0.633** | **0.385** | **0.716** | **0.479** | **0.836** | 0.529 | **0.777** | 0.488 |

### 5.3 Temporal localization

When outputs are collapsed onto the time axis, the student achieves the highest frame AP on NIPS4Bplus, exceeding BirdCODE by 0.120, and matches BirdCODE on XC-AJ (0.836 for both; Table 2), despite BirdCODE being trained specifically for onset-offset detection. Temporal IoU is more mixed: released YOLO11l leads on XC-AJ at 0.556, while BirdCODE leads on NIPS4Bplus at 0.545. SongMAE scores 0.529 and 0.488, respectively, below both released YOLO models on XC-AJ and below BirdCODE on NIPS4Bplus. Together with the smaller IoU gains in two-dimensional localization, this suggests that the student ranks vocalization regions well but that a single threshold calibrated on Powdermill transfers poorly. Collapsing the 2D mask across frequencies further widens this gap, since activation at any frequency extends the predicted interval and merges neighboring events.

## 6. CONCLUSION

We present a pipeline in which a vision-language model annotates avian vocalizations on spectrograms and a bioacoustic encoder student learns from these annotations to localize vocalizations in time and frequency. Trained on fewer than 7 hours of teacher-labeled recordings, the student outperforms the teacher on the development set and outperforms supervised detectors trained on human annotations on held-out time-frequency localization. YOLO11 detectors trained on our teacher labels match those trained on human annotations, suggesting that VLM annotation can substitute for costly expert labeling for this task. Future work could improve annotation through stronger VLMs, better prompts, priors on birdsong structure (e.g., frequency ranges of bird vocalization), improved review agent pipelines, and targeted human corrections.

## 7. COMPLIANCE WITH ETHICAL STANDARDS

This study used only open-access recordings (Xeno-Canto, Powdermill, WABAD, XC-AJ, NIPS4Bplus, Hawaii), no animals were handled, so no ethical approval was required.

## 8. Acknowledgments

The authors have no relevant financial or nonfinancial interests to disclose. The authors used ChatGPT and Claude solely to rephrase, simplify, and improve the manuscript’s prose and grammar, and to assist with programming. All suggestions were reviewed and approved by the authors prior to submission.

## Footnotes

1 https://github.com/ggml-org/llama.cpp

## References

[1] S. Kahl, C. M. Wood, M. Eibl, and H. Klinck, “BirdNET: A deep learning solution for avian diversity monitoring,” Ecological Informatics, vol. 61, pp. 101236, 2021.

[2] D. Stowell, M. D. Wood, H. Pamuła, Y. Stylianou, andH. Glotin, “Automatic acoustic detection of birds through deep learning: The first Bird Audio Detection challenge,” Methods in Ecology and Evolution, vol. 10, no. 3, pp. 368–380, 2019.

[3] G. Vengrovski, M. R. Hulsey-Vincent, M. A. Bemrose, and T. J. Gardner, “TweetyBERT: Automated parsing of bird-song through self-supervised machine learning,” Patterns, vol. 7, no. 4, pp. 101491, 2026.

[4] K. G. Kelly et al., “Estimating population size for California spotted owls and barred owls across the Sierra Nevada ecosystem with bioacoustics,” Ecological Indicators, vol. 154, pp. 110851, 2023.

[5] B. M. Van Doren, A. Farnsworth, K. Stone, D. M. Osterhaus, J. Drucker, and G. Van Horn, “Nighthawk: Acoustic monitoring of nocturnal bird migration in the Americas,” Methods in Ecology and Evolution, vol. 15, no. 2, pp. 329–344, 2024.

[6] M. R. Hulsey-Vincent, G. Vengrovski, E. Sova, and T. J. Gardner, “Lesions involving medial Anterior Forebrain Pathway circuitry destabilize phrase timing in adult canary song,” bioRxiv, p. 2026.07.11.737998, 2026.

[7] W. Vellinga and R. Planqué, “The Xeno-canto collection and its relation to sound recognition and classification,” in Conference and Labs of the Evaluation Forum, 2015.

[8] F. Michaud, J. Sueur, M. Le Cesne, and S. Haupert, “Un-supervised classification to improve the quality of a bird song recording dataset,” Ecological Informatics, vol. 74, pp. 101952, 2023.

[9] A. Fine et al., “BirdCODE: Detecting bird communication at scale,” bioRxiv, p. 2026.07.31.742086, 2026.

[10] S. Hexeberg, F. Tong, H. Vishnu, and M. Chitre, “Time-frequency localization of bird calls in dense soundscapes,” arXiv:2606.10407 [cs.SD], 2026.

[11] M. Hagiwara, “AVES: Animal vocalization encoder based on self-supervision,” in Proc. IEEE Int. Conf. Acoust., Speech, Signal Process. (ICASSP), 2023, pp. 1–5.

[12] L. Rauch, R. Heinrich, I. Moummad, A. Joly, B. Sick, and C. Scholz, “Can masked autoencoders also listen to birds?,”Transactions on Machine Learning Research, 2025.

[13] Y. Zhang and J. Li, “BirdSoundsDenoising: Deep visual audio denoising for bird sounds,” arXiv:2210.10196 [cs.SD], 2022.

[14] J. Redmon, S. Divvala, R. Girshick, and A. Farhadi, “You only look once: Unified, real-time object detection,” in Proc. IEEE Conf. Comput. Vis. Pattern Recognit. (CVPR), 2016, pp. 779–788.

[15] G. Jocher and J. Qiu, “Ultralytics YOLO11,” https://github.com/ultralytics/ultralytics, 2024, Version 11.0.0.

[16] S. Dixit, L. M. Heller, and C. Donahue, “Vision language models are few-shot audio spectrogram classifiers,” arXiv:2411.12058 [cs.SD], 2024.

[17] Qwen Team, “Qwen3.8-27B,” Hugging Face model card, https://huggingface.co/Qwen/Qwen3. 8-27B, 2026, Accessed Sep. 14, 2026.

[18] G. Vengrovski and T. J. Gardner, “SongMAE: A bioacoustic encoder for birdsong,” bioRxiv, p. 2026.08.17.745361, 2026.

[19] I. Loshchilov and F. Hutter, “Decoupled weight decay regularization,” in International Conference on Learning Representations, 2019.

[20] L. Rauch et al., “BirdSet: A large-scale dataset for audio classification in avian bioacoustics,” in The Thirteenth International Conference on Learning Representations, 2025.

[21] L. M. Chronister, T. A. Rhinehart, A. Place, and J. Kitzes, “An annotated set of audio recordings of Eastern North American birds containing frequency, time, and species information,” Ecology, vol. 102, no. 6, pp. e03329, 2021.

[22] L. Jeantet and E. Dufourq, “Improving deep learning acoustic classifiers with contextual information for wildlife monitoring,” Ecological Informatics, vol. 77, pp. 102256, 2023.

[23] C. Pérez-Granados et al., “WABAD: A world annotated bird acoustic dataset for passive acoustic monitoring,” Ecology, vol. 107, no. 2, pp. e70317, 2026.

[24] V. Morfi, Y. Bas, H. Pamuła, H. Glotin, and D. Stowell, “NIPS4Bplus: a richly annotated birdsong audio dataset,” PeerJ Computer Science, vol. 5, pp. e223, 2019.

[25] A. Navine, S. Kahl, A. Tanimoto-Johnson, H. Klinck, and P. Hart, “A collection of fully-annotated soundscape recordings from the Island of Hawai’i,” Zenodo [Dataset], 2022.

